# HNF4α regulates acyl chain remodeling and ether lipid accumulation in hepatic steatosis

**DOI:** 10.1101/2023.06.08.544272

**Authors:** Helaina Von Bank, Gisela Geoghegan, Raghav Jain, Manasi Kotulkar, Mae Hurtado-Thiele, Paula Gonzalez, Charlie Kirsh, Autumn Chevalier, Ian Huck, Kathryn Scheuler, Alan Attie, Mark Keller, Udayan Apte, Judith Simcox

## Abstract

Hepatocyte nuclear factor 4α (HNF4α) is an established transcriptional master regulator of differentiation, maintenance, and metabolism. Polymorphisms in HNF4α are linked to several diseases in humans including diabetes and nonalcoholic fatty liver disease (NAFLD). Identifying novel regulation of lipid metabolism by HNF4α would inform on NAFLD development and progression. We directly assessed HNF4α activity through chromatin immunoprecipitation (ChIP)-sequencing and integration of untargeted lipidomics. Direct regulation by HNF4α can be difficult to assess due to the role of HNF4α in liver homeostasis; to rapidly disrupt activity, mice were exposed to cold stress which induces hepatic steatosis in several hours. Cold exposure shifted HNF4α activity with differential genome occupancy of more than 50% of HNF4α binding sites. Focusing on HNF4α binding to promoter with active transcription determined that HNF4α directly regulates fatty acid desaturation, ether lipid synthesis, and peroxisomal biogenesis in response to cold exposure. Integration of lipidomics found that cold exposure increases the very long chain polyunsaturated fatty acid composition of the hepatic lipid pool, including ether lipids, in an HNF4α dependent manner. Because portions of ether lipid synthesis are in the peroxisome and peroxisomal biogenesis is directly HNF4α regulated, we analyzed peroxisomal abundance and found increases with cold exposure that are ablated with loss of HNF4α. This peroxisomal regulation was independent of PPARα— a known regulator of peroxisomes and lipid metabolism—since loss of HNF4α was not rescued by PPARα overexpression. These data determined that regulation of hepatic steatosis by HNF4α is more complex than triglyceride accumulation and includes acyl chain modifications, ether lipid synthesis, and peroxisomal oxidation.

## Introduction

Hepatocyte nuclear factor 4α (HNF4α) is a master regulator of differentiation and maintenance; deletion of HNF4α leads to hepatocyte dedifferentiation, an inability to metabolize drugs, and decreased hepatocyte proliferation (1–4). HNF4α functions as a homodimer through direct DNA binding to DR1-containing consensus sequences, and HNF4α DNA occupancy is dynamically altered throughout hepatocyte differentiation (5, 6). Beyond its requirement in hepatocyte differentiation, HNF4α also has an established role in regulating glucose and lipid metabolism in mature hepatocytes. Loss of function mutations of HNF4α are known to cause maturity onset diabetes of the young type 1 (MODY1) leading to aberrant glucose regulation and lower circulating lipoprotein A, vLDL, HDL, and triglycerides (7–9). Similar genetic ablation of HNF4α in mice leads to hepatic steatosis, non-alcoholic fatty liver disease (NAFLD), and hepatocellular carcinoma (1, 10–12). Given the numerous functions of HNF4α in cellular identity and lipid metabolism, downstream modulation of specific HNF4α targets that affect hepatic steatosis holds the highest therapeutic potential for NAFLD without side effects.

An acute stress that disrupts homeostasis and alters HNF4α genome occupancy, has the potential to identify HNF4α targets that contribute to hepatic steatosis. Our previous work found that HNF4α was essential for thermogenesis and that acute cold exposure altered lipid processing transcripts controlled by HNF4α (13, 14). We showed that HNF4α directly regulates hepatic acylcarnitine production and liver-specific knockout of HNF4α (HNF4α LKO) results in an inability to produce circulating acylcarnitines and maintain body temperature (15, 16). Because we observed transient changes in acylcarnitine synthesis transcripts regulated by HNF4α with acute cold exposure, we wanted to utilize this model to explore other lipid processing pathways that could be modulated by HNF4α in the liver.

To identify lipid processing pathways regulated by HNF4α during cold exposure, we combined genome occupancy analysis and untargeted lipidomics. Through chromatin immunoprecipitation (ChIP)-sequencing of HNF4α from the livers of cold exposed mice, we observed that HNF4α regulates very long chain polyunsaturated fatty acids and ether lipid synthesis. Combining HNF4α occupancy with active transcription marker Rbp1-CTD determined that HNF4α regulated fatty acid remodeling through the previously observed *Elovl2* and *Scd1,* as well as novel regulators including *Elovl5* and *Fads1* (17–19). These numerous regulators of fatty acid remodeling suggests that HNF4α enables orchestration of a rapid shift that is reflected in the total abundance of very long chain polyunsaturated fatty acids across lipid classes. Similarly, examination of ether lipid synthesis and degradation transcripts in cold exposure identified HNF4α regulation of *Tmem189*, *Pla2g6,* and *Tmem86b.* Liver specific loss of HNF4α led to decreases in ether lipid levels and loss of cold exposure induction of ether lipids and peroxisomes. In cultured hepatocytes, HNF4α regulation of peroxisomal abundance was independent of PPARα, since overexpression of PPARα failed to rescue peroxisomal abundance or ether lipid levels. Together these data strongly suggest an underappreciated role of HNF4α activity in coordinated remodeling of fatty acids, ether lipids, and peroxisomal abundance as well the importance of disrupting metabolic homeostasis to uncover novel regulatory pathways.

## Results

### Loss of HNF4α leads to hepatic steatosis and lower levels of unsaturated fatty acids

HNF4α is an established regulator of hepatic steatosis and NAFLD in humans and mouse models, and genetic ablation of HNF4α alone is sufficient to drive triglyceride accumulation in the liver (1, 10, 20). While accumulation of triglycerides in the liver is detrimental, the manifestation of NAFLD is impacted by acyl chain composition and low abundance lipid species. For example, distinct acyl chain composition and species-specific changes to the lipid pool are known to regulate inflammation, insulin resistance, and fibrosis (21–23). We aimed to identify novel regulation of acyl chain composition and lipid processing pathways by HNF4α. To explore this regulation, we induced liver-specific HNF4α knockout (HNF4α LKO) through adenoviral delivery of Cre under a liver-specific promoter (AAV8-TBG-Cre) in HNF4α^F/F^ mice. One week post AAV8-TBG-Cre administration, we observed that the loss of HNF4α was sufficient to increase liver weight, increase liver triglycerides, and increase liver cholesterol levels (**Figure 1A-D**). While total lipid abundance increased, untargeted liquid chromatography-mass spectrometry (LC-MS) lipidomics revealed dynamic changes in individual lipid species, with 83 lipids significantly increased and 148 decreased in the KO livers (FDR<0.15) (**Figure 1E**). These changes in individual lipid species reflected an overall shift in the saturation of the lipid pool, with a decrease in polyunsaturated acyl chains in HNF4α LKO (**Figure 1F**).

**Figure 1.**
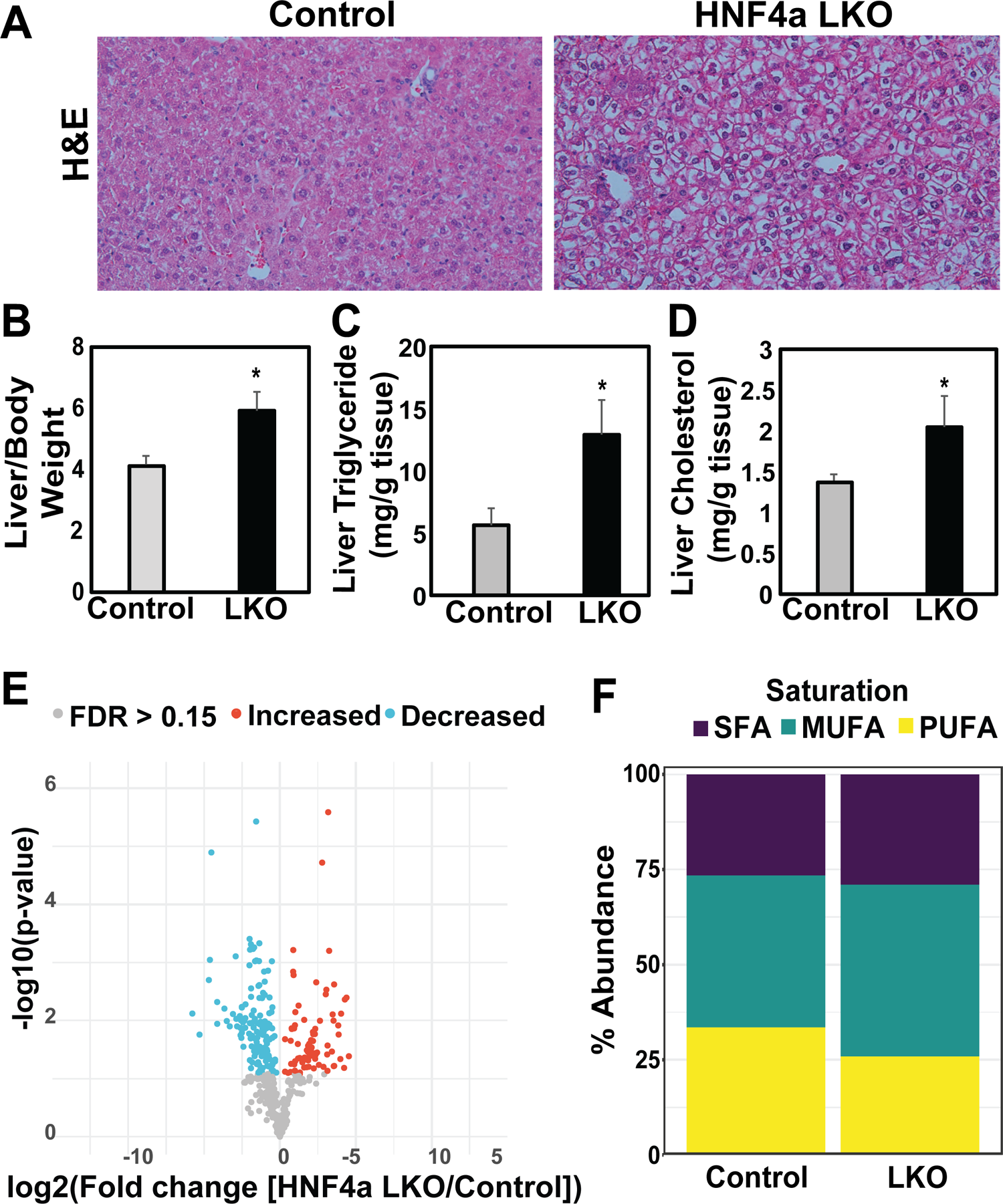
Loss of HNF4α leads to hepatic steatosis and lower levels of unsaturated fatty acids. (A) Representative images of hematoxylin and eosin (H&E) stain of liver sections from HNF4α F/F mice injected with AAV8-TBG-GFP (Control) or AAV8-TBG-Cre (HNF4α LKO). (B) Normalized liver weight, (C) liver triglycerides measured by colorimetric assay, and (D) liver cholesterol measured by colorimetric assay in control or LKO mice (n=3/group). Data are shown as means ± SD; *p<0.05 by unpaired Student’s t-test (E) Volcano plot of lipids detected by untargeted LC/MS lipidomics comparing HNF4α LKO versus control livers. Significantly increased and decreased lipids are colored with red and blue, respectively, with an adjusted p-value (FDR) of 0.15 used as a cutoff. n=3/group. (F) Relative abundance of saturated fatty acids (SFA), monounsaturated fatty acids (MUFA), or polyunsaturated fatty acids (PUFA) in the acyl chains across all lipids detected in the lipidomics analysis.

### Cold exposure exacerbates HNF4α-induced hepatic steatosis and lipid remodeling

HNF4α is known to be important for hepatocyte maintenance and homeostasis. To identify novel lipid regulatory pathways, we aimed to disrupt hepatic lipid homeostasis acutely through cold exposure. We subjected HNF4α LKO mice to cold exposure, a physiological stress that rapidly induces hepatic lipid processing to support thermogenesis. We observed that cold exposure for 5 hours induced triglyceride accumulation in control mice, which was further exacerbated in HNF4α LKO mice (**Figure 2A**). Using publicly available data sets, we explored if cold exposure altered HN4α regulation of lipid metabolism. By integrating RNA-seq data on 850 DO mice to generate a list of correlated transcripts that are driven by HN4α and combined this list with RNA-seq from the livers of mice exposed to room temperature or cold as well as mice that had thermogenic activation with β3-adrenegic receptor activation by CL-316,243 (**Figure S1B &C**). These global shifts in transcripts associated with HN4α expression led us to broadly assess HN4α activity in cold exposure. HNF4α LKO livers showed altered expression of genes associated with hepatic steatosis, including increased expression of *Fasn* with cold exposure as well as decreased expression of *Apoa4* (**Figure S2A**).

**Figure 2.**
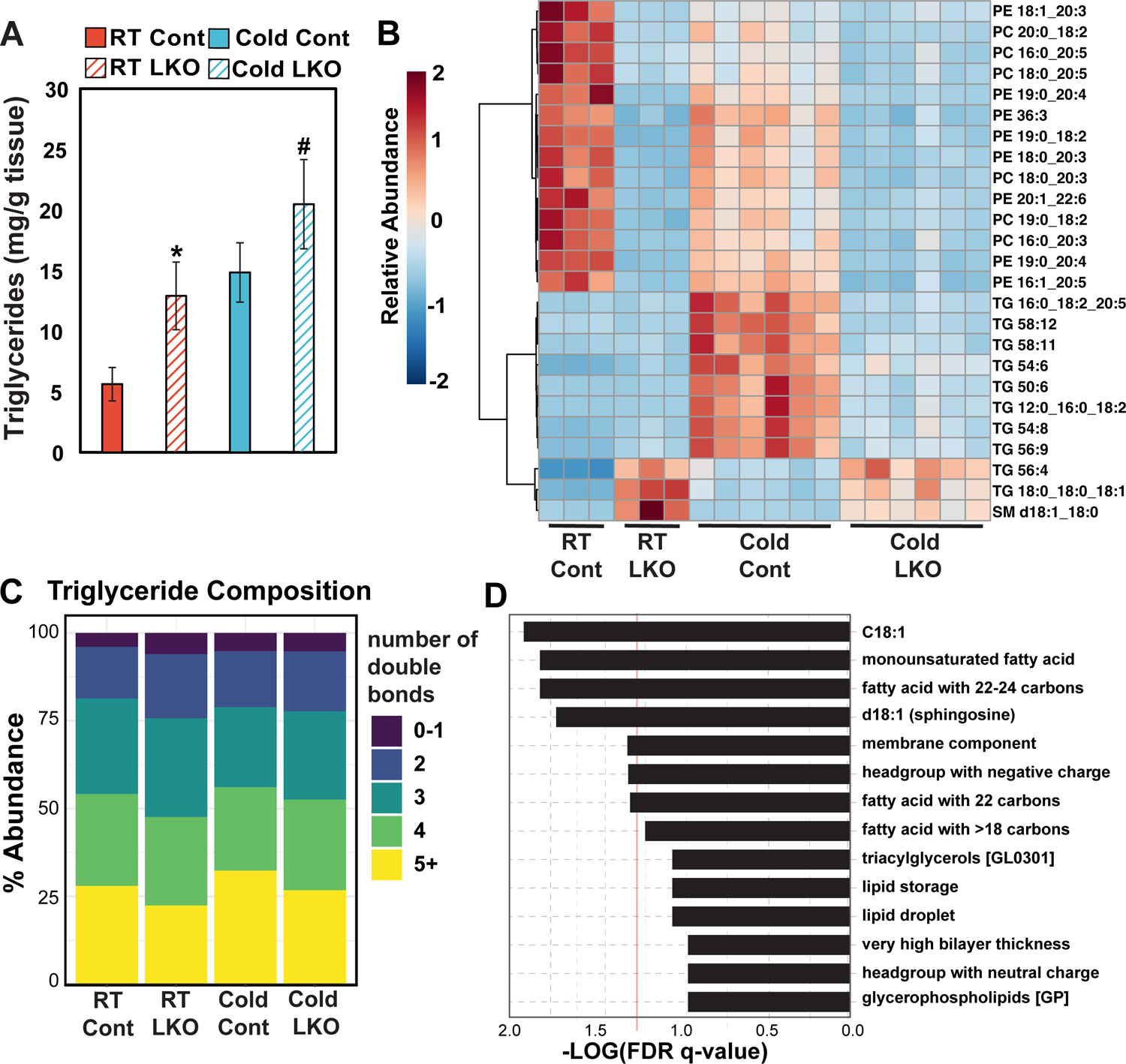
Cold exposure exacerbates HNF4α-induced hepatic steatosis and lipid remodeling. Lipid analyses from livers of control or HNF4α liver-specific knockout (LKO) mice exposed to room temperature (23°C) or cold (4°C) for 6 hours. n = 3-6/group. (A) Total triglycerides measured by colorimetric assay, normalized to tissue weight. Data are shown as means ± SD; significance assessed by ANOVA with post-hoc Tukey HSD comparisons *p<0.05 RT vs Cold Control, #p<0.05 RT vs Cold LKO (B) Cluster analysis (Ward’s method) of the 25 most variable lipids measured by untargeted LC/MS lipidomics. (C) The relative degree of desaturation of triglycerides detected the lipidomics. (D) Lipid ontology (LION) analysis of lipidomics from cold-exposed control or LKO liver. For enrichment analysis, lipid abundances were normalized as a percentage and then ranked based on one-tailed t-test.

We then performed untargeted lipidomics on these livers to investigate species specific lipid changes underlying the hepatic steatosis induced by HNF4α LKO during cold exposure. We observed that several triglyceride species were increased with HNF4α deficiency—such as TG 18:0_18:0_18:1—while many triglycerides containing polyunsaturated acyl chains were induced with cold exposure in control mice but not HNF4α LKO mice. (**Figure 2B**). Analyzing the sum of acyl chains across all detected triglycerides indicated that HNF4α deficiency reduced the degree of desaturation across the triglyceride pool (**Figure 2C**). We also observed phospholipids containing polyunsaturated acyl chains were decreased in HNF4α LKO, suggesting that HNF4α regulates fatty acid processing pathways such as elongation and desaturation. Pathway analysis using lipid ontology (LION) showed that HNF4α LKO combined with cold exposure altered specific acyl chains, including oleic acid (18:1), monounsaturated fatty acids, and fatty acids with 22-24 carbons (24) (**Figure 2D**).

### HNFα ChIP-sequencing with cold exposure identifies novel lipid regulatory targets

Our previous work suggests that HNF4α activity modulates acylcarnitine synthesis during cold exposure (15). To determine changes in global gene occupancy of HNF4α that associated with changes in lipid composition, we performed ChIP-sequencing of HNF4α in the livers of both cold and room temperature exposed mice. We observed that HNF4α DNA occupancy shifted dramatically in response to cold exposure. Differential binding analysis (Diffbind) showed a loss of binding in 13075 sites with just 236 sites gained (**Figure 3A**). These changes in HNF4α occupancy likely reflect altered transcriptional activation, as the active transcription marker Rbp1-CTD showed differential binding, with almost double the number of sites lost than gained (7,357 versus 3,867). H3K27Ac, which labels active enhancers, also lost 10,527 sites while only gaining 4,257 (**Figure 3A**) (25). Changes in HNF4α occupancy occurred mostly in the promoter regions up to 3 kb from the transcriptional start site (**Figure 3B**). Analysis of HNF4α occupancy in the promoter regions showed that many of the binding sites were unique to either room temperature (1508) or cold (917), with a smaller portion shared between the two conditions (997) (**Figure 3C**). Many of these promoter binding sites are functionally characterized to be involved in lipid transport, lipolysis, and fatty acid remodeling (**Figure 3C**). Motif analysis further supported regulation of lipid metabolism since other transcriptional regulators of lipid processing such as PPARα, FOXA2, and ERRα were represented in the binding sites (**Figure 3D**). Gene ontology (GO) of the promoter regions in room temperature exposed livers showed a broad range of hepatocyte maintenance and lipid signaling pathways, including various oxylipids (**Figure S3A**). In contrast, GO analysis of HNF4α promoter occupancy in cold exposure identified pathways associated with lipid remodeling and catabolism such as β-oxidation of very long chain fatty acids, omega-3 and omega-6 metabolism, mitochondrial fatty acid oxidation, and peroxisomal lipid metabolism (**Figure 3E**). Similarly, GO analysis of H3K27ac and Rpb1-CTD gained sites shows enrichment in β-oxidation of long chain fatty acids, omega-3 and omega-6 metabolism, and peroxisomal lipid metabolism (**Figure 3A, Figure S3B-D**). These changes in global H3K27Ac, Rpb1-CTD, and HNF4α occupancy demonstrate that there is a dynamic transcriptional response to cold exposure directed to specific lipid metabolic pathways.

**Figure 3.**
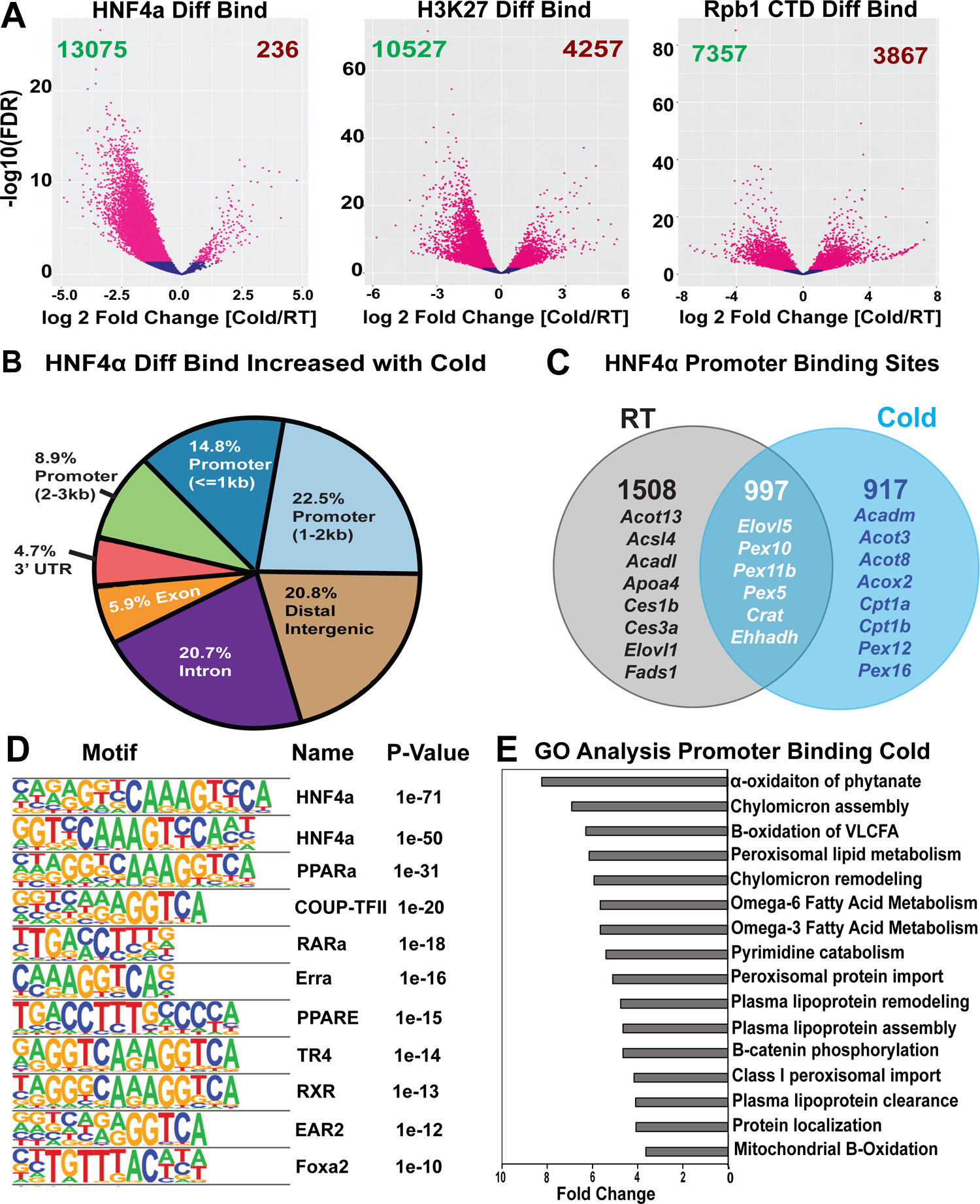
HNFα ChIP-sequencing with cold exposure identifies novel lipid regulatory targets. (A) Differential binding site analysis of HNF4α, H3K27ac and Rpb1-CTD occupancy in cold exposure versus room temperature. (B) Annotation of HNF4α binding sites that are gained in cold exposure. (C) Overlap of HNF4α binding sites in the promoter region as defined by <3 kb of the start site between room temp and cold with select genes highlighted. Black indicates unique promoter occupancy in room temp, blue indicates promoter occupancy in cold exposure, and white indicates occupancy in both room temp and cold. (D) Motif analysis of known binding sites within HNF4α ChIP DNA. (E) Gene ontology (GO) analysis of the promoter binding sites in cold exposure. Biological replicates for ChIP-seq n=2.

### Hnf4α regulates cold-induced hepatic fatty acid elongation and desaturation

Cold exposure induced HNF4α occupancy of omega-3 and omega-6 lipid metabolism promoters (**Figure 3E**), correlating with the observed decreased abundance of polyunsaturated acyl chains in lipids from HNF4α LKO livers with cold (**Figure 2C**). To validate if this change in binding associated with the omega fatty acid composition of the lipid pool, we directly assessed the acyl chains from all lipids detected by LC-MS lipidomics and observed increased incorporation of essential fatty acids (18:2, 18:3) as well as their modified products (20:4, 20:5, 22:4, 22:5, 22:6) with cold exposure (**Figure 4A**). Omega fatty acids—such as the omega-6 linoleic acid (18:2)—are modified by desaturases that add double bonds and elongases that increase the chain length in two-carbon increments (**Figure 4B**). Direct assessment of the relative abundance of omega-6 acyl chains showed a decreased ratio of arachidonic acid (20:4) to linoleic acid (18:2) in HNF4α LKO livers relative to controls, indicating that HNF4α deficiency disrupts the omega-6 modification pathway (**Figure 4C)**.

**Figure 4.**
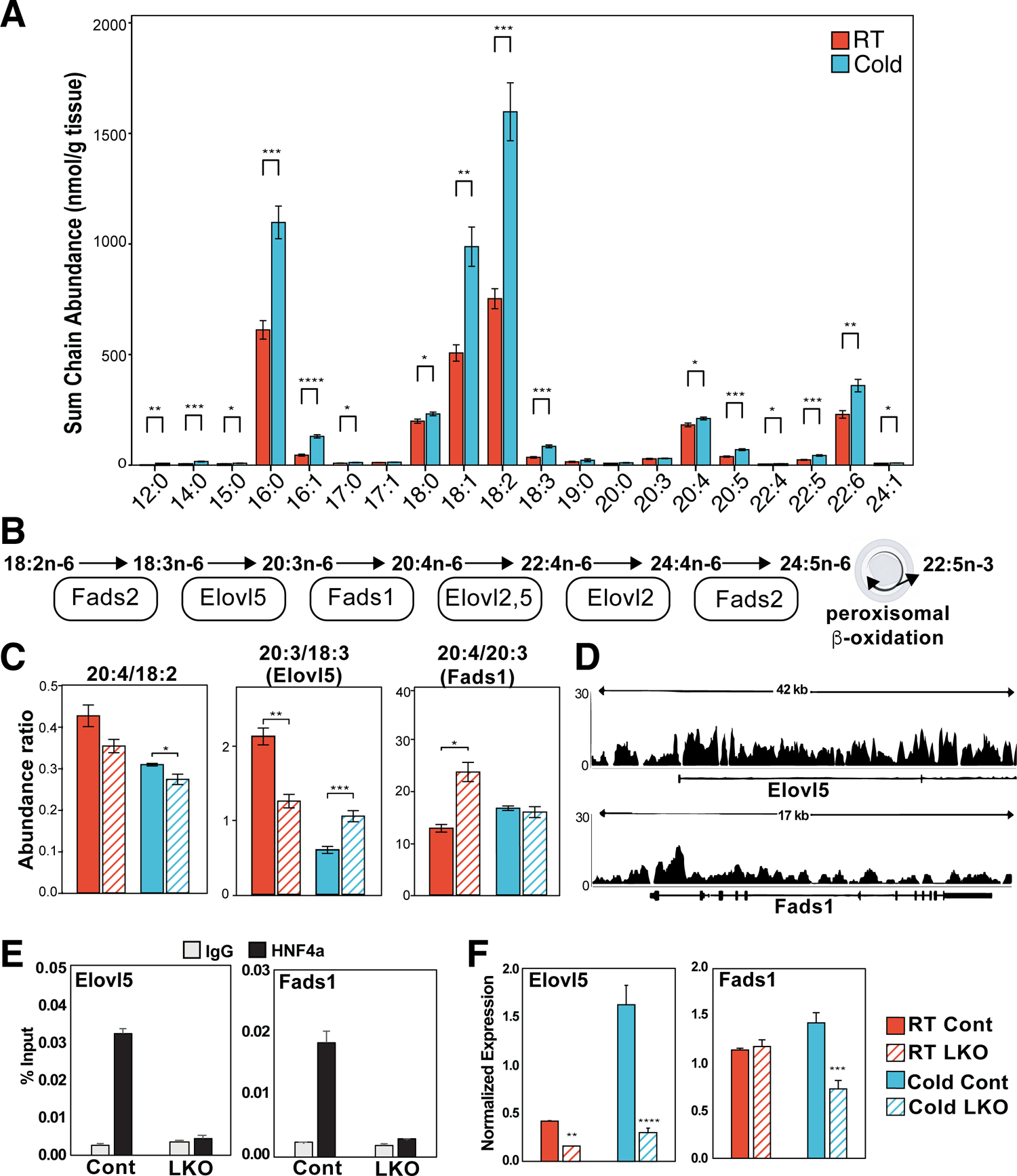
Hnf4α regulates cold-induced hepatic fatty acid elongation and desaturation. (A) The sum of different acyl chains from all lipids detected by LC-MS lipidomics of liver from room temperature or cold exposed mice, from Jain et al., 2022 (14). n=6/temperature group (B) Schematic of the omega-6 fatty acid processing pathway. (C) Ratios of the sum abundance of specific acyl chains across all lipids detected by LC-MS lipidomics of control and liver-specific HNF4α knockout (LKO) livers. Ratios that correspond to the activity of Elovl5 and Fads1 are noted. n=3-6/group. Data are shown as means ± SD; significance assessed by ANOVA with post-hoc Tukey HSD comparisons *p<0.05; **p<0.01; ***p<0.001. (D) HNF4α occupancy tracks on elongation and fatty acid desaturation genes in the liver in cold exposed mice. (E) ChIP-qPCR of HNF4α in livers of control and HNF4α LKO mice. n=3 per group. (F) Gene expression analysis of livers from control and liver specific HNF4α knockout mice, n=3-6 per group.

These lipid desaturation and elongation changes were associated with HNF4α occupancy of the enzymes that regulate these fatty acid modificaitons. We observed HNF4α ChIP tracks in pathway genes (*Elovl5*, *Elovl2*, *Fasd1*, *Fads2*) and confirmed direct binding in the promoters of *Elovl5*, *Fads1*, and *Fads2* by targeted ChIP qPCR (**Figure 4D&E, Figure S4B&C**). Cold exposure increased the expression of *Elovl5*, *Fads1*, and *Fads2*, reflecting the increased abundance of omega-derived acyl chains in the hepatic lipid pool (**Figure 4 A&F, Figure S4D**). While HNF4α deficiency reduced expression of *Elovl5* and *Fads2* independent of temperature condition, expression of *Fads1* was reduced by HNF4a deficiency specifically in cold exposure **(Figure 4F)**. Beyond the omega fatty acid pathway, we also observed regulation of other fat modifying genes by HNF4α, including the saturated fatty acid pathway. *Elovl6*, which catalyzes the initial elongation of palmitate (16:0) to oleate (18:0), was increased with HNF4α deficiency, while the downstream elongase *Elovl3* had completely ablated expression in HNF4α LKO (**Figure S4D**). These data show that HNF4α directly regulates acyl chain diversity through elongation and desaturation of fatty acids.

### Ether lipid acyl chains are particularly impacted by loss of HNF4α

From our untargeted lipidomics analysis, we observed that HNF4α deficiency altered the levels of many ether lipids in both room temperature and cold conditions (**Figure 5A**). Ether glycerophospholipids such as plasmalogens contain unique ether bonds at the sn1 position, giving them specific functions in membrane dynamics, cell signaling, and oxidative stress (26). We observed that most ether lipids were decreased by HNF4α deficiency, especially in the room temperature condition. Because ether lipids often contain a polyunsaturated acyl chain, their decrease in HNF4α LKO livers could be due to disrupted omega fatty acid processing. However, we hypothesized that the disruption of ether lipid levels across many species may also be due to regulation of their biosynthesis or degradation. Ether lipid synthesis initiates in the peroxisome, yet none of the peroxisomal genes (*Gnpat, Agps, Far1*) had significantly changed expression in HNF4α LKO livers (**Figure S5A)**. Ether lipid intermediates produced by the peroxisome are then processed by the ER into mature ether lipids, including plasmalogens produced by Tmem189 (**Figure 5B**) (27). *Tmem189* expression was significantly reduced in HNF4α LKO livers with both temperature condition and cold exposure (**Figure 5C**). This reduction in expression was likely due to direct HNF4α activity, as we observed HNF4α ChIP tracks in the *Tmem189* gene and binding in the promoters by ChIP-qPCR (**Figure 5 D&E**). Plasmalogen levels can also be regulated by degradation pathways that hydrolyze plasmalogens and lysoplasamogens (**Figure 5B**). The phospholipase and putative plasmalogenase *Pla2g6* had increased expression with HNF4α deficiency, while the lysoplasmalogenase *Tmem86b* had decreased expression that associated with direct HNF4α binding activity (**Figure 5C-E**). Other potential regulators of ether lipids such as *Pmpx4*—which has been associated with levels of hepatic ether lipids— also showed HNF4α-dependent expression and direct HNF4α binding (**Figure S5A-C**)(28). These data indicate that HNF4α regulates ether lipid levels at multiple points, including polyunsaturated substrate availability, biosynthesis, and degradation.

**Figure 5.**
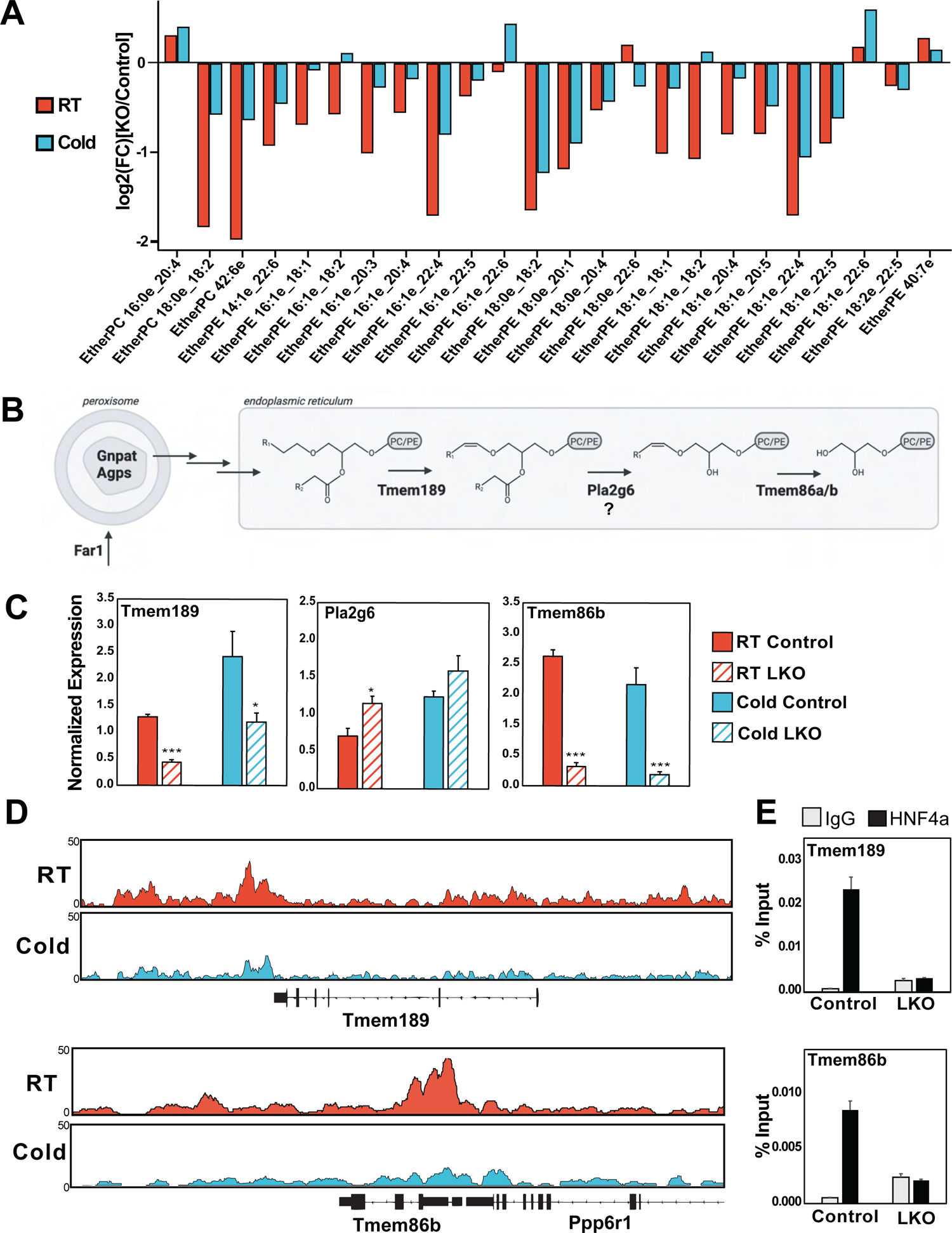
Ether lipid acyl chains are particularly impacted by loss of HNF4α. (A) Log2 fold change of ether lipids between HNF4α liver-specific knockout (LKO) and control livers at room temperature or cold detected by global lipidomics analysis. n=3-6 per group. (B) Schematic of the plasmalogen synthesis and degradation pathway with known and putative proteins annotated. (C) Gene expression analysis of plasmalogen synthesis and degradation genes in livers of control and HNF4α LKO mice, n=3-6 per group. (D) HNF4α occupancy tracks of *Tmem189* and *Tmem86b* in the liver of mice exposed to room temperature or cold. (E) HNF4α ChIP-qPCR for *Tmem189* and *Tmem86b* in livers of control and HNF4α LKO mice. n=3 per group. Data are shown as means ± SD; significance assessed by ANOVA with post-hoc Tukey HSD comparisons *p<0.05; **p<0.01; ***p<0.001.

### Peroxisome abundance is impacted by HNF4α regulation

Beyond direct regulation of biosynthesis and degradation genes, changes in ether lipids may also be driven by decreased peroxisomal biogenesis. GO analysis from our ChIP-seq indicated that HNF4α binding to several peroxisomal pathways was enriched in cold exposure (**Figure 3E**). This binding activity associated with increased peroxisomal abundance in the liver during cold exposure, as measured by immunofluorescent staining of Pmp70, a VLCFA transporter localized to the peroxisome membrane (**Figure 6A&B**). Cold exposure induced expression of the Pmp70 gene (*Abcd3),* as well as the peroxisomal biogenesis factor *Pex16,* in an HNF4α-dependent manner (**Figure 6E**). We confirmed direct binding of HNF4α to *Abcd3* and *Pex16* (**Figure 6C&D**), suggesting that HNF4α induces peroxisomal biogenesis and activity during cold exposure. Similar regulation was observed in other peroxisomal biogenesis regulators including *Pex5, Pex11a,* and *Pex14* as well as peroxisomal lipid processing enzymes *Acox1* and *Ehhadh* (**Figure S6A-E**). We then wanted to test if this regulation required peroxisome proliferator-activated receptor α (PPARα), a well-established activator of peroxisomal metabolism that is also transcriptionally regulated by HNF4α. To pursue this, we performed siRNA knockdown of HNF4α along with lentiviral overexpression of PPARα in the murine hepatocyte Hepa1-6 cell line. Knockdown of HNF4α decreased PMP70 protein levels, which was not rescued with lentiviral overexpression of PPARα **(Figure 6F)**. Similarly, gene expression of *PMP70*, *Pex16*, and *Tmem189* was not rescued by overexpression of PPARα **(Figure 6F and Figure S6F)**. These data suggest that hepatic peroxisomes are induced during cold exposure and HNF4α is directly regulating peroxisomal abundance.

**Figure 6.**
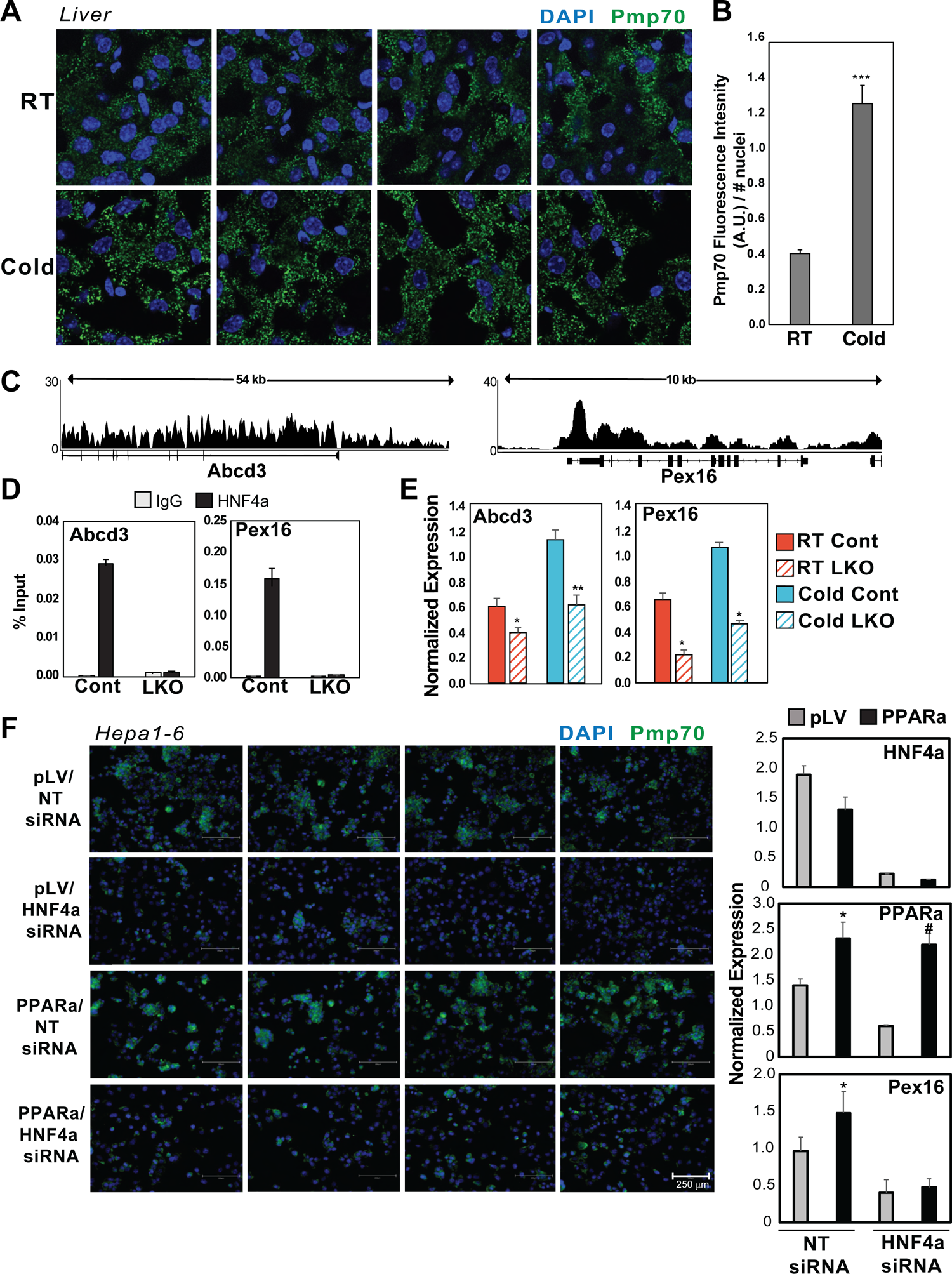
Peroxisome abundance is impacted by HNF4α regulation. (A) Immunofluorescence staining of Pmp70 (Alexa Fluor488) and nuclei (DAPI) of room temperature (top) and cold exposed livers (bottom) along with (B) quantification of Pmp70 signal relative to number of nuclei. (C) HNF4α occupancy tracks of *Abcd3* and *Pex16* in the liver. (D) HNF4α ChIP-qPCR of *Abcd3* and *Pex16* in livers of control and liver-specific HNF4α knockout (LKO) mice. n=3 per group. (E) Gene expression of *Abcd3* and *Pex16* in livers of control and HNF4α LKO mice, n=3-6 per group. (F) Immunofluorescent staining of Pmp70 in Hepa 1-6 cells with either empty vector (pLV) or PPARa overexpression along with non-targeting (NT) siRNA or HNF4α siRNA and corresponding gene expression of *HNF4α*, *PPARα*, and *Pex16*. n=4 per group Data are shown as means ± SD; significance assessed by ANOVA with post-hoc Tukey HSD comparisons *p<0.05; **p<0.01; ***p<0.001.

## Discussion

HNF4α is an established regulator of plasma bile acids, cholesterol, and lipoprotein particles and directly contributes to the development of NAFLD. We wanted to identify lipid processing pathways regulated by HNF4α that contribute to the development of hepatic steatosis, allowing for future targeting independent of HNF4α function in hepatocyte maintenance. To disrupt HNF4α regulation of lipid metabolism acutely, we used cold exposure which drives liver lipid accumulation in four hours. By combining ChIP-seq with untargeted lipidomics, we determined that cold exposure rapidly remodels HNF4α DNA occupancy to regulate fatty acid processing pathways, ether lipid levels, and peroxisomal biogenesis.

Our data demonstrate that HNF4α remodels the fatty acid pool through coordinated desaturation and elongation. We observed that very long chain polyunsaturated fatty acyl chain abundance was dependent on HNF4α since HNF4α LKO lowered 20:4, 20:5, and 22:6 acyl chains, and that the ratio of omega-6 pathway product arachidonic acid to its starting substrate linoleic acid (20:4/18:2) was decreased. Interestingly, expression of *Fads1,* which directly catalyzes the production of arachidonic acid, was increased specifically in the cold in a HNF4α-dependent manner. We also found changes in saturated fatty acid processing, including HNF4α-dependent increases in the ratios of 18:0/16:0 that correlates with expression of *Elovl6* (**Figure S4**). However, evaluation of acyl chain ratios is limited as they do not represent levels of the FFAs themselves and may not reflect total flux through these pathways, which have enzymes acting on multiple substrates. While expression of *Elovl5* was decreased with HNF4α LKO, we did not see a consistent decrease in the ratio of 20:3/18:3, potentially due to other factors that regulate the diversity of acyl chain substrates such as mitochondrial and peroxisomal oxidation or uptake of exogenous lipids. Others have previously shown that *Elovl2* is a direct target of HNF4α and that HNF4α LKO leads to decreased expression of *Fads2* and several *Elovl* genes (17, 29). Here we build upon these finding by confirming regulation of the entire elongation and desaturation cascade and reveal potentially specific regulation of *Fads1* and acyl chain remodeling during cold exposure.

We observed direct HNF4α regulation of genes involved in peroxisomal biogenesis and peroxisomal oxidation. This regulation led to changes in peroxisomal abundance and ether lipid abundance in HNF4α LKO during cold exposure (**Figure 5&6**). These results are surprising given that most control of peroxisomes by HNF4α has been attributed to its regulation of PPARα (30). It is likely the activity or perhaps the cooperation of both transcription factors is needed to promote cold induced peroxisomal biogenesis since motif analysis of HNF4α binding sites showed enrichment for the PPAR motif (**Figure 3D**). More work is needed to understand the interplay of regulation between PPARα and HNF4α, since our work suggests that PPARα overexpression is not sufficient to rescue the lower peroxisomal abundance caused by HNF4α knockdown in cultured hepatocytes. Recent work has demonstrated that a double knockout of HNF4α and PPARα defended against hepatic steatosis with increased 18:1/18:0 fatty acid ratios (29). Future studies are needed to explore coregulator binding, and regulatory complex formation at shared sites of regulation.

It is unclear why peroxisomes and β-oxidation are induced in the cold-exposed liver. Acox1 is the rate limiting enzyme and first step in peroxisomal β-oxidation and is strongly induced in cold exposure (**Figure S6**). The first step of peroxisomal β-oxidation generates FADH2 but since the peroxisome doesn’t have a respiratory chain to produce ATP, electrons are instead transferred to O2 generating heat (26, 31). The liver has been shown to contribute to body temperature through heat production, but the mechanism has been unknown (32). Perhaps peroxisomal abundance and peroxisomal β-oxidation is increased during cold exposure to generate heat and aid in thermogenesis.

Surprisingly, cold exposure completely alters HNF4α genome occupancy causing a loss of 13075 binding sites and a gain of 236 sites (**Figure 3**). The mechanisms that control the cold-induced HNF4α genome occupancy remain unknown, but two potential mechanisms include co-regulator binding and ligand binding. Co-regulators such as PGC-1α, p300, SMRT, and GRIP1 have all been shown to bind HNF4α to regulate hepatocyte and intestinal differentiation and maintenance programs (33). The binding of these co-regulators to HNF4α in acute metabolic stresses such as cold exposure has not been explored. Differential ligand binding could also alter HNF4α activity since endogenous ligand linoleic acid binding is altered in the fasted vs fed state (34). Whether altered ligand binding also regulates HNF4α activity in cold exposure has not been explored but may be an important mechanism as cold-induced hepatic lipid remodeling program depends on adipose tissue lipolysis, similar to the fasted state (15). One other possibility is altered HNF4α isoform abundance with cold exposure, since various pairs of isoform dimers differentially regulate gene programs such as inflammation (6). More work is needed to identify mechanisms of HNF4α regulation, and cold exposure holds promising potential in determining basic physiological regulation.

Our findings that HNF4α is a regulator of acute lipid remodeling in cold exposure has implications in numerous metabolic diseases including fatty liver disease. There are several hundred GWAS studies that identify HNF4α polymorphisms as being associated with cholesterol levels, type 2 diabetes, cardiovascular disease, and bile acid production (35–38). Using acute cold exposure to push lipolysis and hepatic lipid processing, we have further characterized HNF4α’s regulation of lipid processing pathways such as elongation and desaturation as well as uncovered novel HNF4α regulation of peroxisomal biogenesis and peroxisomal fatty acid oxidation. The impact of HNF4α’s role in the global lipid regulatory program has numerous physiological states to be explored including disease progression of type 2 diabetes and hyperlipidemia.

## STAR Methods

| REAGENT or RESOURCE | SOURCE | IDENTIFIER |
| --- | --- | --- |
| Antibodies |  |  |
| rabbit anti-HNF4α antibody | Abcam | ab181604 |
| rabbit anti-Acetyl-Histone H3 (Lys27) (D5E4) | Cell Signaling | 8173 |
| mouse anti-Rpb1 CTD | Cell Signaling | 2629 |
| Chemicals, peptides, and recombinant proteins |  |  |
| SPLASH Lipidomix Mass Spec Standard | Avanti Polar Lipids | Cat#330707-1EA |
| Dulbecco's Phosphate-Buffered Saline (DPBS) | Thermo Fisher | Cat#14190144 |
| Methanol | Sigma-Aldrich | Cat#1060351000 |
| Methyl tert-butyl ether | Fisher Chemical | Cat#1634-04-4 |
| Isopropanol | Honeywell | Cat#34965 |
| Water | Honeywell | CAS 7732-18-5 |
| Acetonitrile | Honeywell | Cat#34967 |
| Formic acid | Thermo Fisher | Cat#A117 |
| Ammonium formate | Thermo Fisher | Cat#A115-50 |
| Formulab standard chow | Lad Diet | Cat#5008 |
| Trizol Reagent | Thermo Fisher | Cat#15596018 |
| Critical commercial assays |  |  |
| SimpleChIP Plus Enzymatic Chromatin IP Kit | Cell Signaling | 9005 |
| ChIP DNA was purified using Zymo ChIP DNA Clean and Concentrator Kit | VWR | D5205 |
| Qubit® dsDNA HS Assay Kit | Invitrogen | Q32851 |
| Deposited data |  |  |
| ChIP-seq of HNF4α from room temperature and cold exposure | This paper | GEO: |
| Lipidomics on HNF4α control and LKO livers in room temperature and cold | This paper | Metabolights: MTLBS4474 |
| R Code | This paper | hcvonbank GitHub |
| Experimental models: Cell lines |  |  |
| Hepa 1-6 Cells | ATCC | CRL-1830 |
| Experimental models: Organisms/strains |  |  |
| C57BL6/J Mice, male, 12 weeks old | Jackson Laboratories | Stock#000664 |
| HNF4α F/F Mice, Male, 12 weeks old | Huck et al. 2019 | N/A |
| Oligonucleotides |  |  |
| siRNA targeting sequence: HNF4α | Horizon Discovery | L-065463-00-0005 |
| Primers for RT-PCR and targeted ChIP, see Table S1 and S2 | This paper ordered from IDT | N/A |
| Recombinant DNA |  |  |
| AAV8-TBG-GFP | Vector Biolabs | VB1724 |
| AAV8-TBG-Cre | Vector Biolabs | VB1743 |
| Software and algorithms |  |  |
| ImageJ | Schneider et al., 2012 | https://imagej.nih.gov/ij/ |
| Genome Scans Software | Linke et al., 2020 | N/A |
| Agilent MassHunter Lipid Annotator Version 10 | Koelmel *et al.*, 2020 | N/A |
| Agilent MassHunter Profinder Version 10 | Agilent | N/A |
| Agilent MassHunter Data Acquisition Version 10 | Agilent | N/A |
| Agilent MassHunter Qualitative Analysis Version 10 | Agilent | N/A |
| R Version 4.0.2 | R Core Team | https://www.r-project.org/ |
| QuantStudio Software V1.5.1 (QuantStudio 3 and 5 systems) | Thermo Fisher | https://www.thermofisher.com/us/en/home/global/forms/life-science/quantstudio-3-5-software.html |
| Recombinant DNA |  |  |
| Agilent 6546 Quadrupole Time of Flight LC/MS System | Agilent | N/A |
| Infinity 1290 II UHPLC System | Agilent | N/A |

### Mice

All procedures were conducted in compliance with protocols approved by the University of Wisconsin-Madison and the Kansas University Medical Center Institutional Animal Care and Uses Committees (IACUC). Mice were housed in in groups of 3-5 at 22°C-24°C using a 12 hr light/12 hr dark cycle. Animals were provided ad libitum access to food and water, except when explicitly state for cold tolerance tests. Mice were housed in groups of 2-5 per cage, in standard polypropylene 11.5×7.5×5” containers with enrichment and corncob bedding. Mice were fed a normal chow (Formulab Diet, 5008) diet from the age of 4 weeks. Unless specified in the figure legend, experiments were performed on male C57BL6J male mice. HNF4α F/F mice were male on a C57BL6J background, control mice were HNF4α F/F injected with AAV8-TBG-GFP (Control) while knockouts were HNF4α F/F injected with AAV8-TBG-Cre (HNF4α LKO) one week prior to cold exposure as previously described (15).

### Cold tolerance test

Mice were individually housed in cages with no bedding at 4°C for 6 hours. Food was removed at the T0 start time, zeitgeber time 4, but the mice maintained free access to water. Body temperature was monitored hourly by rectal probe (Physiotemp, RET-3), and mice were checked every 15 minutes for signs of distress or hypothermia. *Correlation Analysis DO* Quantitative trait loci (QTL) mapping involved scanning the genome and testing for association between the 8-state haplotype probabilities and the expression data from the livers of 850 diversity outbred mice (**Supplemental Figure 1**). The genome scans were performed for each gene transcript as previously described (39). This approach determined transcripts that were positively and negatively correlated to HNF4a. A Corr>|0.4| was used as a cutoff. This yielded a subset of 508 genes. Data from RNA-seq of mice exposed to cold or treated with β3-adrenergic receptor agonist (CL-316,243) were incorporated to assess the correlation to HNF4α expression in the DO livers. Transcripts found in the DO mouse lists and the CL-316,243 included 506 genes, while transcripts found in the DO list and cold exposure list matched 502 genes. False discovery rate (FDR) and p-adj values were set at <0.05. The plot shows the respective fold change and correlation to HNF4α expression with the sizes proportional to the FDR/p-adj emphasizing those with larger fold changes.

### ChIP-seq

ChIP was performed on livers from individually housed C57BL/6J mice that were placed at 4°C without food for 6 hours compared to a non-fasted room temperature control group. SimpleChIP Plus Enzymatic Chromatin IP Kit (Cell signaling, 9005) was used according to manufacturer’s instructions. Briefly, 100mg of tissue was used per mouse and three immunoprecipitations were performed with rabbit anti-HNF4α antibody (Abcam, ab181604), rabbit anti-Acetyl-Histone H3 (Lys27) (D5E4) (Cell signaling, 8173), and mouse anti-Rpb1 CTD (4H8) (Cell signaling, 2629). ChIP DNA was purified using Zymo ChIP DNA Clean and Concentrator Kit (VWR, D5205). Purified immunoprecipitated and input DNA was submitted to the University of Wisconsin-Madison Biotechnology Center. DNA concentration and sizing were verified using the Qubit® dsDNA HS Assay Kit (Invitrogen, Carlsbad, California, USA) and Agilent DNA HS chip (Agilent Technologies, Inc., Santa Clara, CA, USA), respectively. Samples were prepared according the TruSeq® ChIP Sample Preparation kit (Illumina Inc., San Diego, California, USA) with minor modifications. Libraries were size selected for an average insert size of 350 bp using SPRI-based bead selection. Quality and quantity of the finished libraries were assessed using an Agilent Tapestation and Qubit® dsDNA HS Assay Kit, respectively. Paired end 150bp sequencing was performed on the NovaSeq 6000. Sequenced reads checked for: sequencing quality, insert length, read concordance, duplications, and adapters via FastP and then aligned to the mouse genome (UCSC build mm10) via bowtie2 v2.2.5 using the sensitive end-to-end read alignment mode. Aligned reads are sorted and indexed with SAMtools 1.9 and peaks are called at <= p0.05 and q0.05 (separately) using the input/IgG as the background control with the MACS2. Primary and Differential Peaks were annotated using annotatePeaks.pl for their positioning relative to genomic elements with Homer v4.11. *De novo* and known motif analysis were carried out with Homer findMotifsGenome.pl. ChIPseeker (v1.28.3) and UCSC.mm10.known gene database in combination with narrowPeak data from MACS to plot. Raw data will be deposited in GEO datasets.

### RNA extract, RT-PCR, RNA-sequencing

RNA was purified using Invitrogen TRIzol reagent (Fisher Scientific, 5596018) according to manufacturer’s instructions. cDNA was synthesized using Invitrogen SuperScript VILO master mix (Fisher Scientific, 11755250). qPCR was performed on an ABI Quant Studio 5 with Applied Biosystem PowerUP SYBR Green Master Mix (Fisher Scientific, A25778). Primers can be found in Supplemental Table 1. For RNA-seq RNeasy mini kit was used (Qiagen, 74106). Samples were submitted to the University of Wisconsin-Madison Biotechnology Center for RNA sequencing. Briefly, RNA size and purity were verified using Qubit® RNA HS Assay Kit (Invitrogen, Carlsbad, California, USA) and Agilent RNA HS chip (Agilent Technologies, Inc., Santa Clara, CA, USA), respectively. Samples were prepared according the TruSeq® Stranded mRNA Library Prep kit (Illumina Inc., San Diego, California, USA) with minor modifications. Libraries were purified using AMPure XP beads (Beckman Coulter, A63880) and sequenced on the NovaSeq 2×150 bp with 30 million reads per sample. Reads were trimmed using trimming software skewer (Jiang et al.2014), filtered for at or near zero expression, and gene expression normalization was carried out by the method of trimmed mean of M-values (TMM). The trimmed single- or paired-end reads are aligned against the selected reference genome sequence using STAR (Spliced Transcripts Alignment to a Reference) (Dobin et al. 2013). Analysis of differentially expressed genes is performed with a glm using the edgeR package (Robinson, McCarthy, and Smyth 2010).

### Correlation Analysis DO

Quantitative trait loci (QTL) mapping involved scanning the genome and testing for association between the 8-state haplotype probabilities and the expression data from the livers of 850 diversity outbred mice (**Supplemental Figure 1**). The genome scans were performed for each gene transcript (39). The software uses linear modeling with sex and DO breeding generation as additive covariates taking random effect into account for the kinship structure of the DO mice. We identified suggestive QTL at LOD > 6.0 and significant QTL at LOD > 7.4. These threshold values were estimated by permutation analysis to obtain a family-wise error correction for genome-wide QTL search. The family-wise error rate ensures that the maximum LOD score across the genome-wide search when applied to a trait with no QTL (i.e., a permuted trait) will exceed the threshold with a fixed probability. For the lenient threshold 6.0, the genome-wide probability of false QTL detection is 0.20. For the stringent threshold 7.4, the genome-wide error rate is controlled at 0.05. The lenient threshold is used to identify the almost-significant associations that co-localize on the genome in hotspots. This approach determined transcripts that were positively and negatively correlated to HNF4α. A Corr>|0.4| was used as a cutoff. This yielded a subset of 508 genes. Data from RNA-seq of mice exposed to cold or treated with CL-316,243 were incorporated to assess the correlation to HNF4α expression in the DO livers. Transcripts found in the DO mouse lists and the CL-316,243 included 506 genes, while transcripts found in the DO list and cold exposure list matched 502 genes. False discovery rate (FDR) and p-adj values were set at <0.05. The plot shows the respective fold change and correlation to HNF4α expression with the sizes proportional to the FDR/p-adj emphasizing those with larger fold changes.

### Western Blot

Protein lysate from liver tissue from C57BL/6J mice (25 mg) or Hepa 1-6 cells were extracted by harvesting in RIPA buffer (Fiher, NC9484499) with protease inhibitor cocktail (Fisher, PIA32953). Homogenization was performed with a Tissue Lyzer II from Qiagen. Lysates were then spun at 4°C for 20 minutes at 14,000 g. Lipid layer aspirated and samples were spun again. Protein concentrations were assayed using Peirce BCA assay (ThermoFisher Scientific, 23227). 20 μg of protein lysates were run on SDS page, transferred onto nitrocellulose (Fisher, #45004003), blocked in 5 % milk in TBS-T and blotted with the following antibodies diluted 1/1000 in 5% milk overnight at 4°C: rabbit anti-HNF4α (R&D Systems, PP-H1415-00), mouse anti-B-Actin Fisher, MAB8929), and rabbit anti-PMP70 (ThermoFisher, PA1-650). Blots were washed with TBS-T and incubated in goat anti mouse secondary antibody (ThermoFisher, A-10668) diluted 1/5000 in 5% milk in TBS-T. at room temp for 1 hour. Blots were washed with TBS-T and developed using Amershan ECL Prime (Fisher, 45002401) and imaged on an iBright C1000.

### siRNA

Hepa 1-6 cells were purchased from ATCC (CRL-1830) and were cultured in DMEM (ThermoFisher,11-965-118) supplemented with 10% FBS (ThermoFisher, 10-437-028) and grown under 5%CO2 at 37°C in accordance with ATCC recommendations for growth. ON-TARGETplus Non-targeting Control Pool (Horizon Discovery, D-001810-10-05) ON-TARGETplus Mouse HNF4a siRNA pool (Horizon Discovery, L-065463-00-0005) were transfected at the concentration of 10 pmol per well in a 12-well dish of Hepa 1-6 cells using RNAiMAX <u>lipofectamine</u> (ThermoFisher, 13778030). Cells were harvested 48 hours after transfection.

### Immunofluorescence

Hepa 1-6 cells were grown on coverslips and washed with PBS 48 hours post transfection with siRNAs. Cells were fixed with 4% EM grade formaldehyde (VWR, 100503-917) in PBS for 15 minutes. Cells were then washed with PBS and blocked in 5% normal goat serum (Sigma, G9023) and permeabilized with 0.1% saponin (Bio Basic, SB4521) in PBS for 1 hour, washed again, and incubated with rabbit anti-PMP70 ((ThermoFisher, PA1-650) diluted 1:1000 in 1%BSA, 0.1% saponin in PBS overnight at 4°C. Cells were washed with PBS and then incubated with goat anti-rabbit Alexa Fluor 488 (ThermoFisher, A11008) diluted 1:1000 in 1%BSA, 0.1% saponin in PBS at room temperature in the dark for 1 hour. Cells were then washed in PBS and coverslips mounted onto slides with VECTASHIELD Antifade mounting medium (Vector Laboratories, H-1000-10) and imaged on Invitrogen EVOS M5000 imaging scope (Fisher, 1256363).

### Lipid Extraction

Liver (25 mg) was homogenized in ceramic 1.4mm bead tubes (Qiagen, Cat. No. 13113-50) containing ice-cold PBS (Gibco #14190144, 250 µL), methanol (215 µL), MTBE (Fisher #1634-04-4, 750 µL) and Splash I Lipidomix internal standard (10 µL) (Avanti, Cat. No. 330707) with a Qiagen TissueLyzer II (Cat. No. 9244420). The sample were then centrifuged at 16,300 g for 5 min at 4 °C. The upper organic phase (500 µL) was transferred to a new tube and evaporated to dryness under vacuum. Samples were resuspended in 150 µL isopropanol and diluted 1:30 in isopropanol for positive mode runs and 1:2 in isopropanol for negative mode runs.

### Colorimetric assays

Total triglycerides were measured from the organic extractions using the L-Type Triglyceride M Assay (Wako) and cholesterol was measured with Amplex Red Cholesterol Assay kit (ThermoFisher #A12216).

### LC/MS analysis

Extracted lipids were separated on an Agilent 1290 Infinity II LC System through a VanGuard BEH C18 guard column (Waters 18003975) then an Acquity C18 column (Waters 186009453, 1.7 μm 2.1 x 100 mm) maintained at 50°C. Injection volumes were 3 µL for positive mode and 5 µL for negative mode. A chromatography gradient was run at a flow rate of 0.500 mL/min composed of mobile phase A (ACN:water [60:40 v/v] in 10 mM ammonium formate and 0.1% formic acid) and mobile phase B (IPA:water [90:10 v/v] in 10 mM ammonium formate and 0.1% formic acid). Both mobile phases were used at varying proportions over 20 minutes to create a gradient with a consistent flow rate of 0.500 mL/min. The chromatography gradient started at 15% mobile phase B increasing to 30% by 2.40 min, 48% by 3.00 min, 82% by 13.20 min, and 99% by 13.80 min. Mobile phase B was then maintained at 99% until 15.40 minutes, then decreased to 15% by 16.00 minutes and held at 15% to 20.00 minutes.

Runs were performed in both positive and negative mode on an Agilent 6546 quadrupole time-of-flight MS dual AJS ESI mass spectrometer connected to the LC system. For both modes, the source gas temperature was set to 325°C, with a gas flow of 12 L/min, nebulizer pressure of 35 psi, sheath gas temperature of 375°C and sheath gas flow of 11 L/min. VCap voltage was set at 4000 V, fragmentor at 190 V, skimmer at 75 V and Octopole RF peak at 750 V. Reference masses were 121.050873 and 922.009798 m/z for positive mode and 112.985588700 and 966.00072500 m/z for negative mode. Samples were acquired in the mass scan range of 119-1500 m/z (positive) and 100-1500 m/z (negative) at a rate of 3 spectra/sec. Tandem mass spectrometry was performed with a fixed collision energy of 25 V at a scan rate of 2 spectra/sec. Raw data is uploaded to Metabolights.

### Data processing and statistics

Results from LC/MS experiments were evaluated using Agilent MassHunter Workstation. Raw data was evaluated using Agilent MassHunter Qualitative Analysis and stored in .d format. Data from six iterative MS/MS analyses of pooled samples were used to generate a library using LipidAnnotator (Agilent) with a mass accuracy of <0.05 ppm. Duplicate annotations with a retention time difference less than 0.1 min were manually removed. Lipid features were extracted and integrated based on the generated library using Profinder Analysis (Agilent) and peak heights were exported for curation and statistical analysis in R (version 4.0.2). Compounds were removed if their abundance in the extraction control was higher that the sample average. Redundant lipid annotations were identified by correlation analysis (R > 0.999) and removed. For volcano plotting, significant lipids were identified by Student’s t-test with a false-discovery rate (FDR) adjustment. Statistical comparison of two groups was done using Student’s t-test. For assessment of ether lipids and acyl chain composition, lipid abundance was normalized to a representative internal standard associated with the head group. Diacylglycerols were excluded to do lack of signal for the internal standard. For lipid ontology (LION) analysis, lipid inputs were normalized as a percentage and ranked by one-tailed T-test p-value. Distributions of associated LION-terms were evaluated by one-tailed Kolmogorov-Smirnov [K-S] test to identify enriched terms (24). Multiple group comparison was assessed by ANOVA with post-hoc Tukey HSD (*p ≤ 0.05, **p ≤ 0.01, ***p ≤ 0.001, ****p ≤ 0.0001).

## Acknowledgements

The authors would like to thank members of the Simcox lab for reviewing and offering critical feedback on the manuscript including Jess Davidson, Gina Wade, Edrees Rashn, and Isabella James as well as Michaela Morhaus. Research reported in this publication was supported by the Glenn Foundation and American Federation for Aging Research to JS (A22068); Hatch Grant to JS and RJ (WIS04000-1024796); JDRF to JS (JDRF201309442); an R01 through NIDDK to JS (R01DK133479), an NSF GRFP to HVB. The content is solely the responsibility of the authors and does not necessarily represent the official views of the National Institutes of Health. The work was also supported in part by startup funds from the University of Wisconsin-Madison School Department of Biochemistry to JS. Figures were made with BioRender.

## Supplemental Figures

Figure S1. (A) Replicate images from H&E staining of livers from control or HNF4α LKO mice. (B) Correlation analysis between expression of HNF4α in the Diversity Outbred (DO) mouse population and associated transcripts that are modulated with cold exposure (blue) and β3-adrenergic receptor agonist CL-316, 243 (green). (C) Gene ontology (GO) of transcripts that are increased with increased HNF4α expression, Cold exposure, and CL-316,243 treatment.

Figure S2 (A) Gene expression analysis of livers from control and liver specific HNF4α knockout mice, n=3-6 per group. n=4 per group Data are shown as means ± SD; significance assessed by ANOVA with post-hoc Tukey HSD comparisons *p<0.05. (B) Representative MS chromatogram from LC-MS lipidomics of control or liver-specific KO livers in positive ionization mode and (C) negative ionization mode. (D) Peak abundance of the triglyceride class internal standard (TG 15:0_18:1(d7)_15:0) across all samples in LC-MS lipidomics analysis. All lipids were normalized to a representative internal standard for that class.

Figure S3. (A) Gene ontology (GO) analysis of HNF4α promoter binding sites (as defined by <3 kb of the start site) in livers from the room temperature condition. (B) GO analysis of HNF4α binding sites gained during cold exposure. (C) GO analysis of Rbp1-CTD binding sites gained during cold exposure. (D) GO analysis of H3K27ac binding sites gained during cold exposure. Biological replicates for ChIP-seq n=2.

Figure S4. (A) Specific acyl chain sums from all lipids detected by LC-MS lipidomics of livers of control or HNF4 α LKO). n=6/ group (B) Schematic of the omega-6 processing pathway. (B) HNF4α occupancy tracks on elongation and fatty acid desaturation genes from in the liver of cold exposed mice. (C) ChIP-qPCR of HNF4α in livers of control and liver specific HNF4α knockout mice. n=3 per group. (D) Gene expression analysis of livers from control and liver specific HNF4α knockout mice, n=3-6 per group. (E) Ratios of the sum abundance of specific acyl chains across all lipids detected by LC-MS lipidomics of control and liver specific HNF4α knockout livers. Ratios that correspond to the activity of Elovl6 and Elovl1/Elovl3/Elovl7 are noted. n=3-6/group. Data are shown as means ± SD; significance assessed by ANOVA with post-hoc Tukey HSD comparisons *p<0.05; **p<0.01; ***p<0.001.

Figure S5. (A) Normalized abundance of ether lipids identified from global lipidomics in the livers of room temperature and cold exposed mice adapted from Jain et al., 2022 (14). (B) Expression of genes related to ether lipid metabolism from livers of control and liver-specific HNF4α knockout (LKO) mice, n=3-6 per group. (C) HNF4α occupancy track of binding to *Pmpx4* in livers from room temperature or cold exposed mice. (D) HNF4α ChIP-qPCR of *Pmpx4* in livers of control and liver specific HNF4α LKKO mice. n=3 per group. Data are shown as means ± SD; significance assessed by ANOVA with post-hoc Tukey HSD comparisons *p<0.05; **p<0.01; ***p<0.001.

Figure S6. (A) Expression of genes related to peroxisomal metabolism from livers of control and liver-specific HNF4α knockout mice, n=3-6 per group. Data are shown as means ± SD; significance assessed by ANOVA with post-hoc Tukey HSD comparisons *p<0.05; **p<0.01; ***p<0.001.(B) HNF4α occupancy track of binding to peroxisomal genes in liver of cold exposed mice. HNF4α ChIP-qPCR of peroxisomal genes from livers of control and liver specific HNF4α knockout mice. n=3 per group. (D) Pmp70 protein expression in the liver of mice exposed to cold along with β-actin expression as a control. (E) HNF4α, Pmp70, and β-actin protein expression in Hepa1-6 cells transfected with either non-targeting (NT) or HNF4α siRNA.

---

**Supplemental Table 1.**
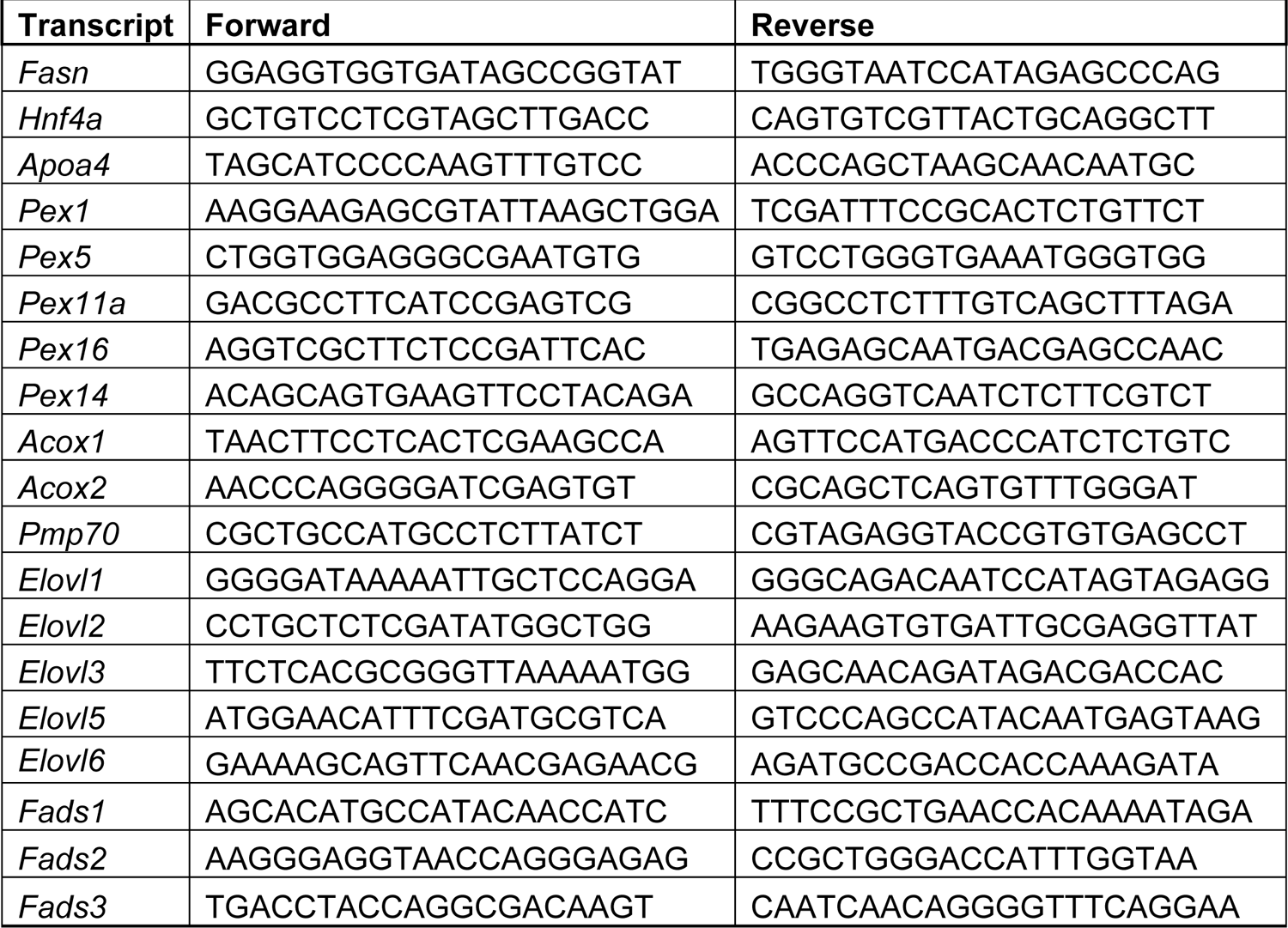
RT-PCR

**Supplemental Table 2.**
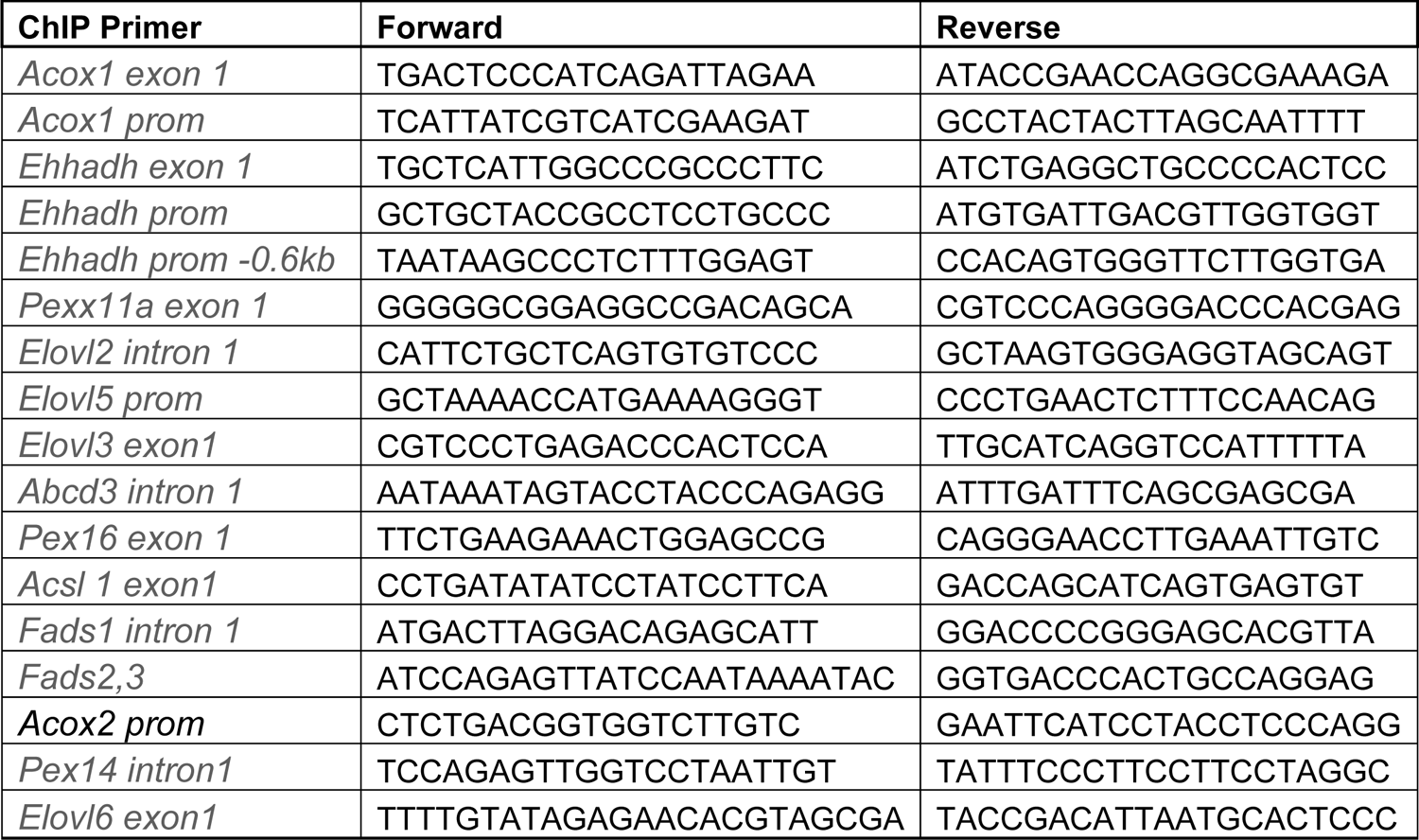
Targeted ChIP Primers

