## Supplementary figures and images for "HNF4α regulates acyl chain remodeling and ether lipid accumulation in hepatic steatosis"

### Supplemental figures

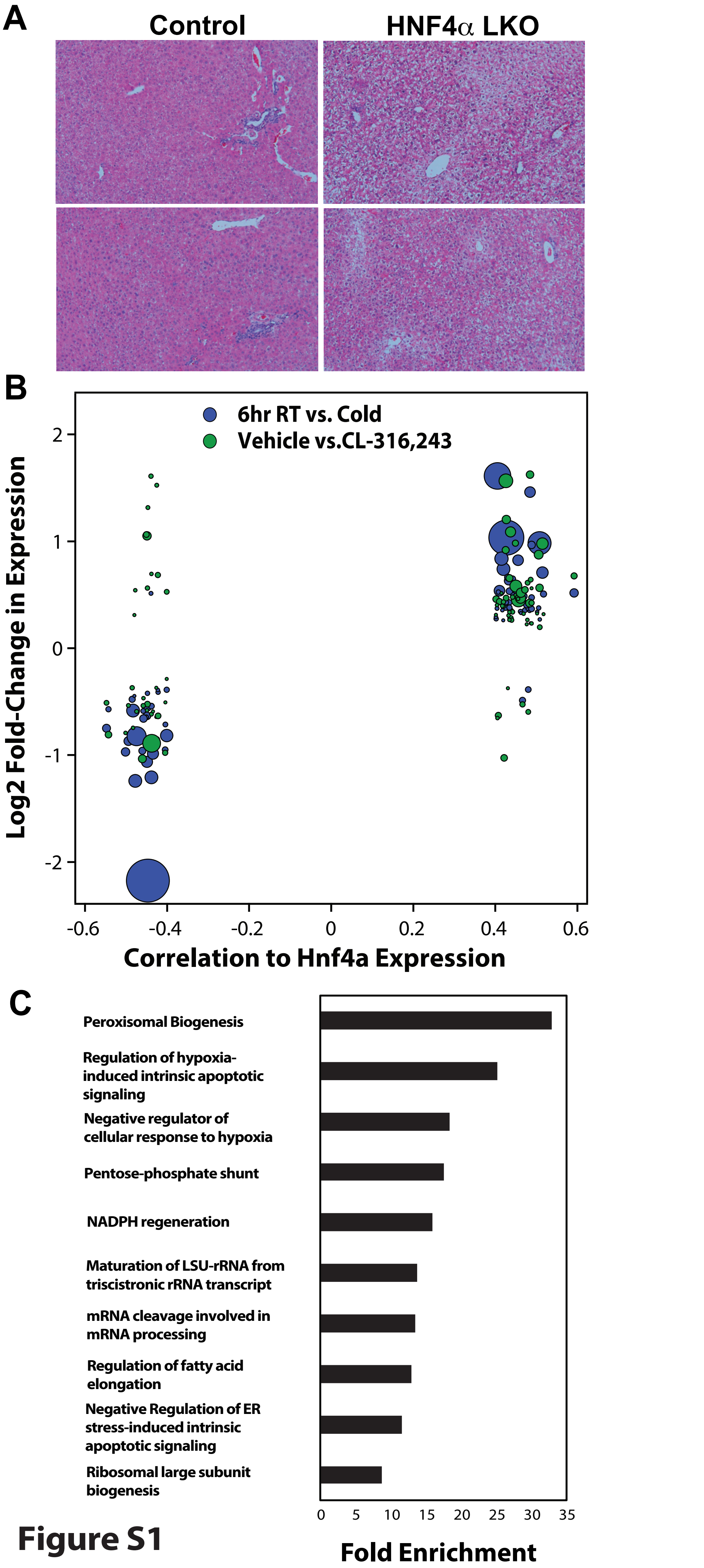

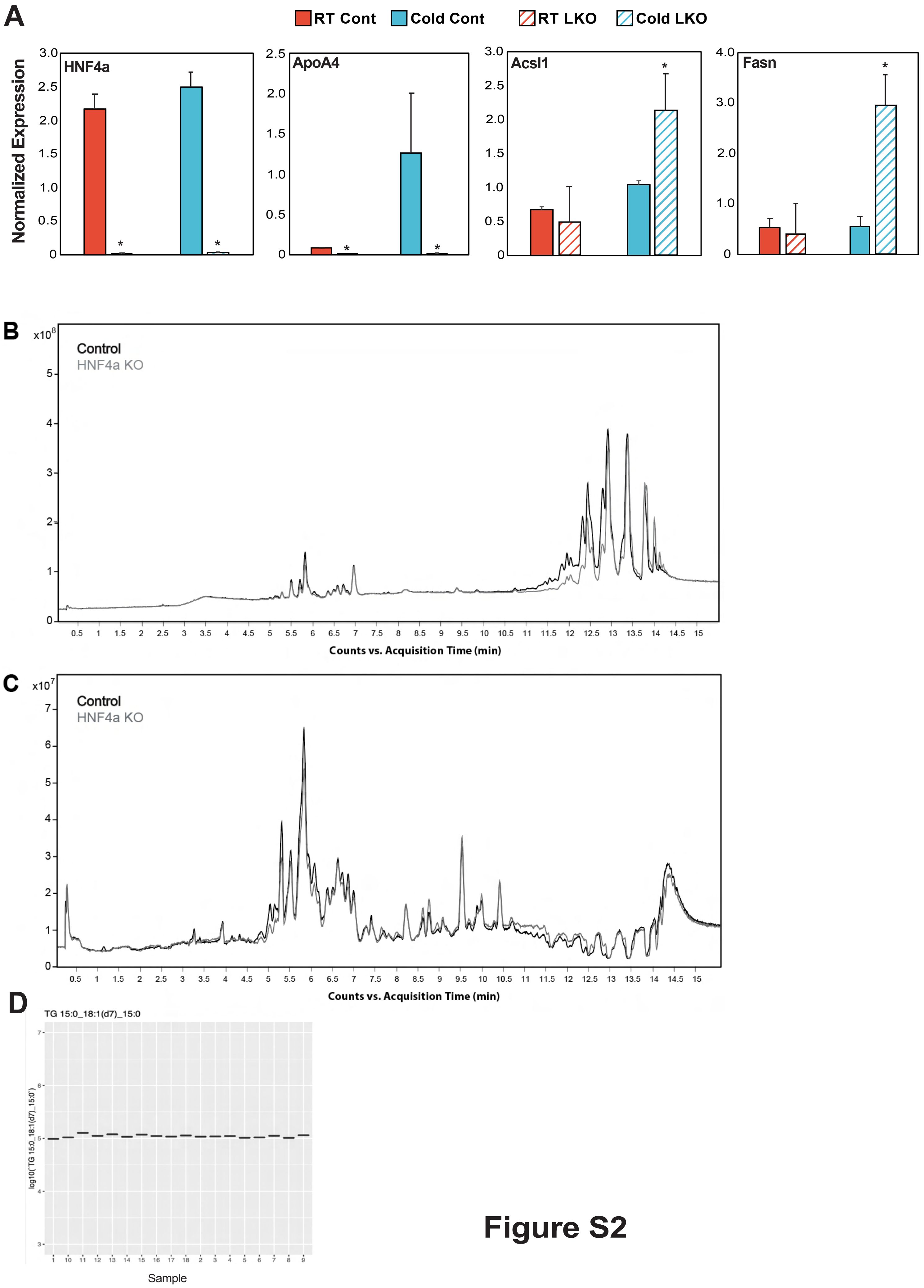

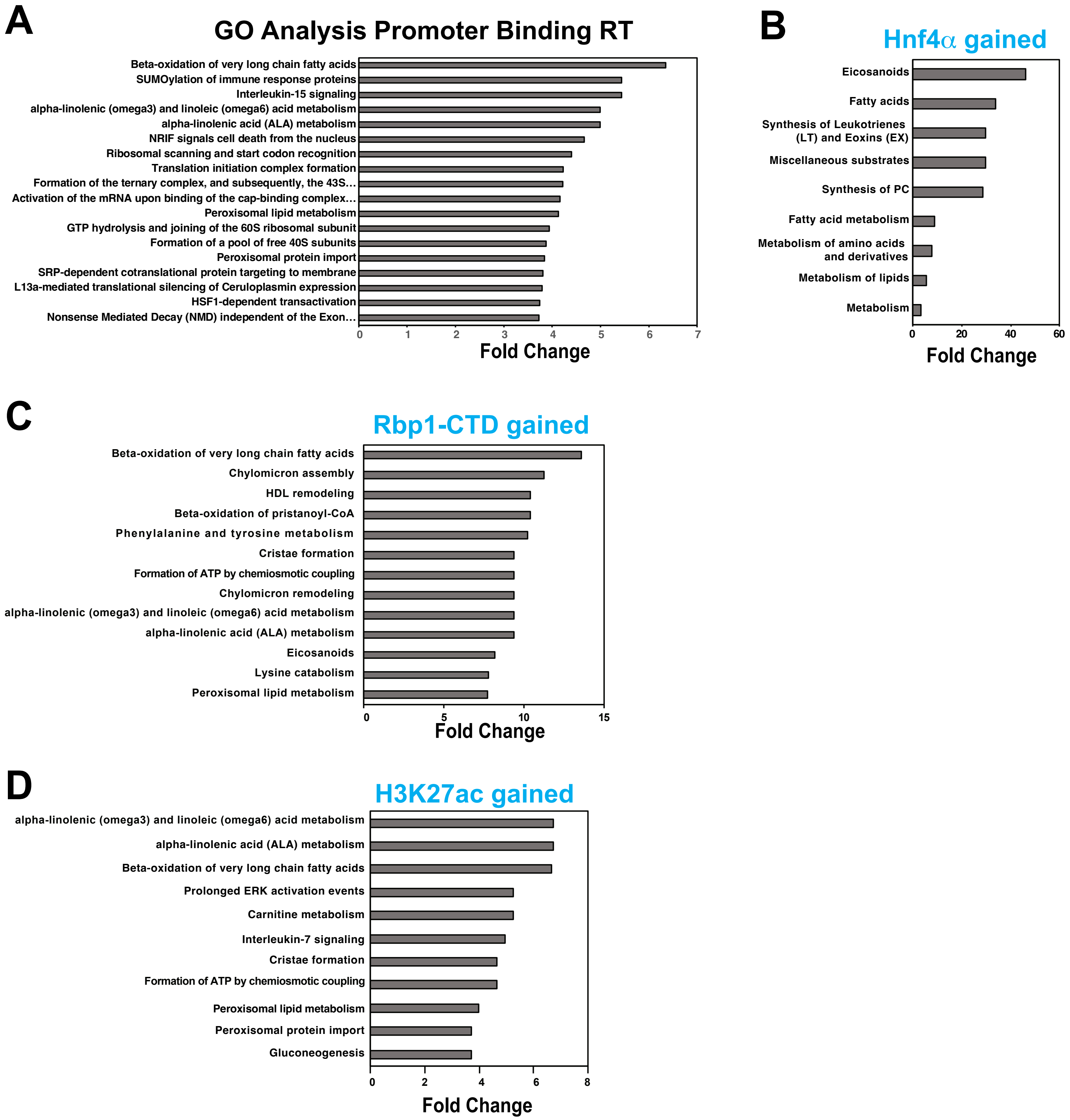

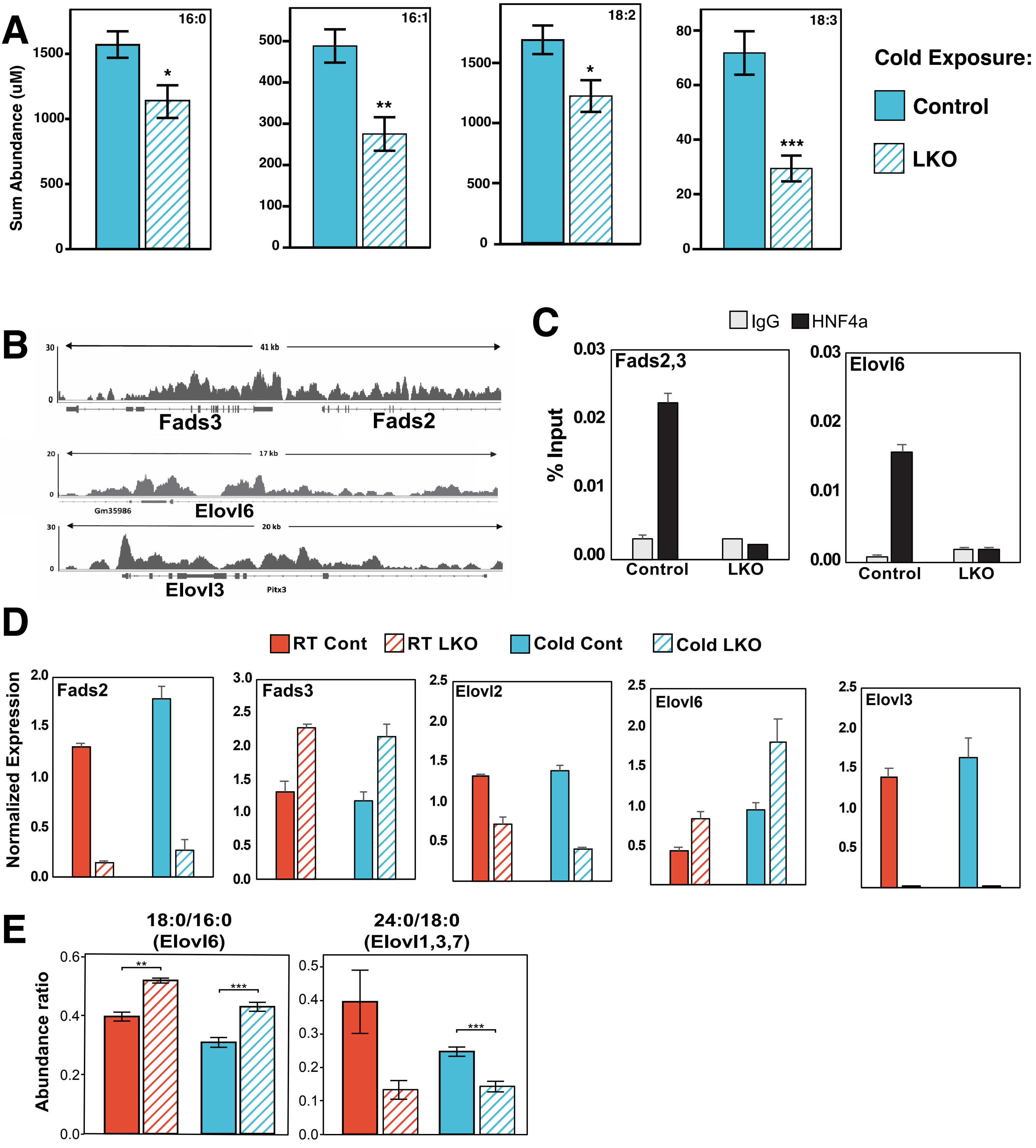

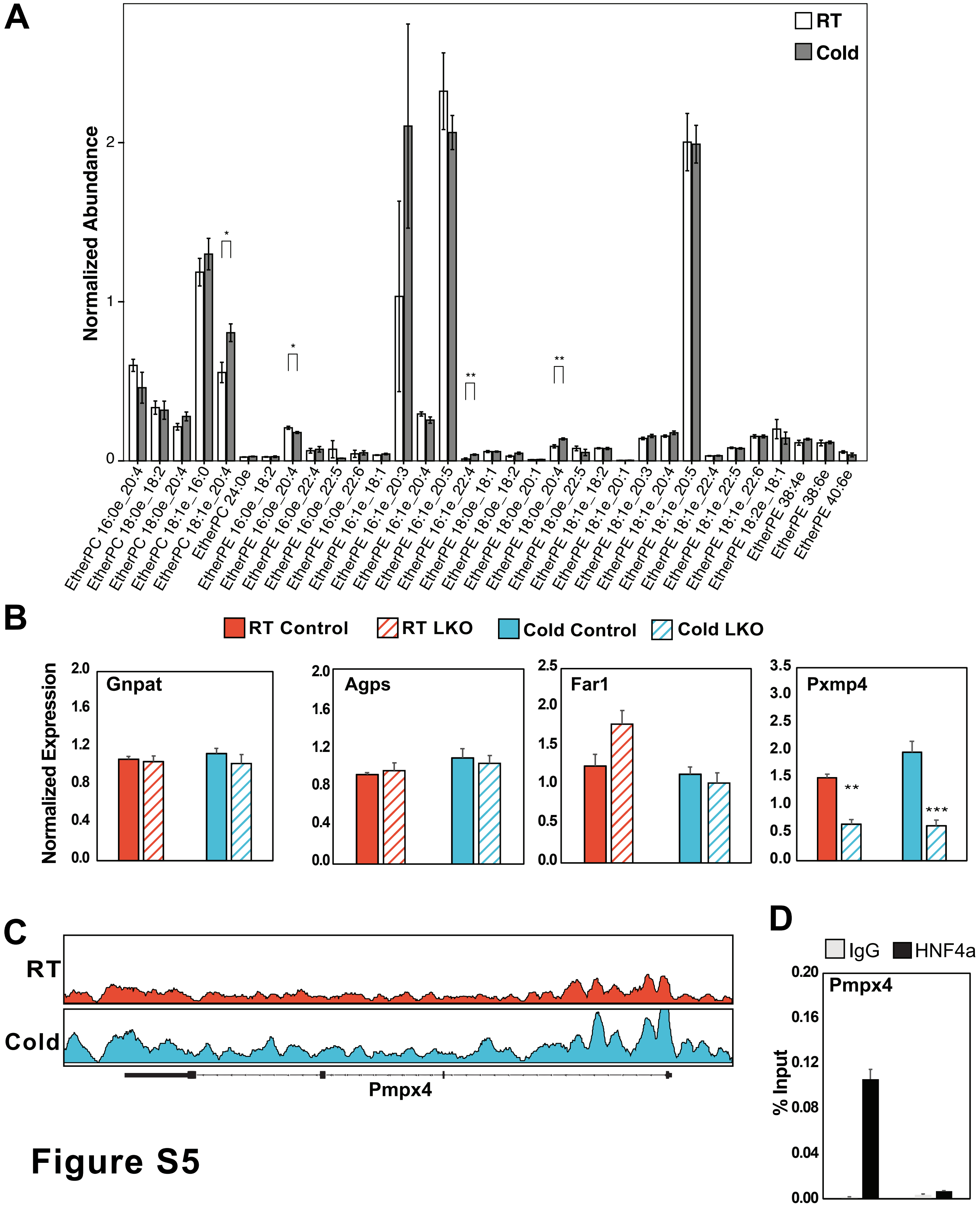

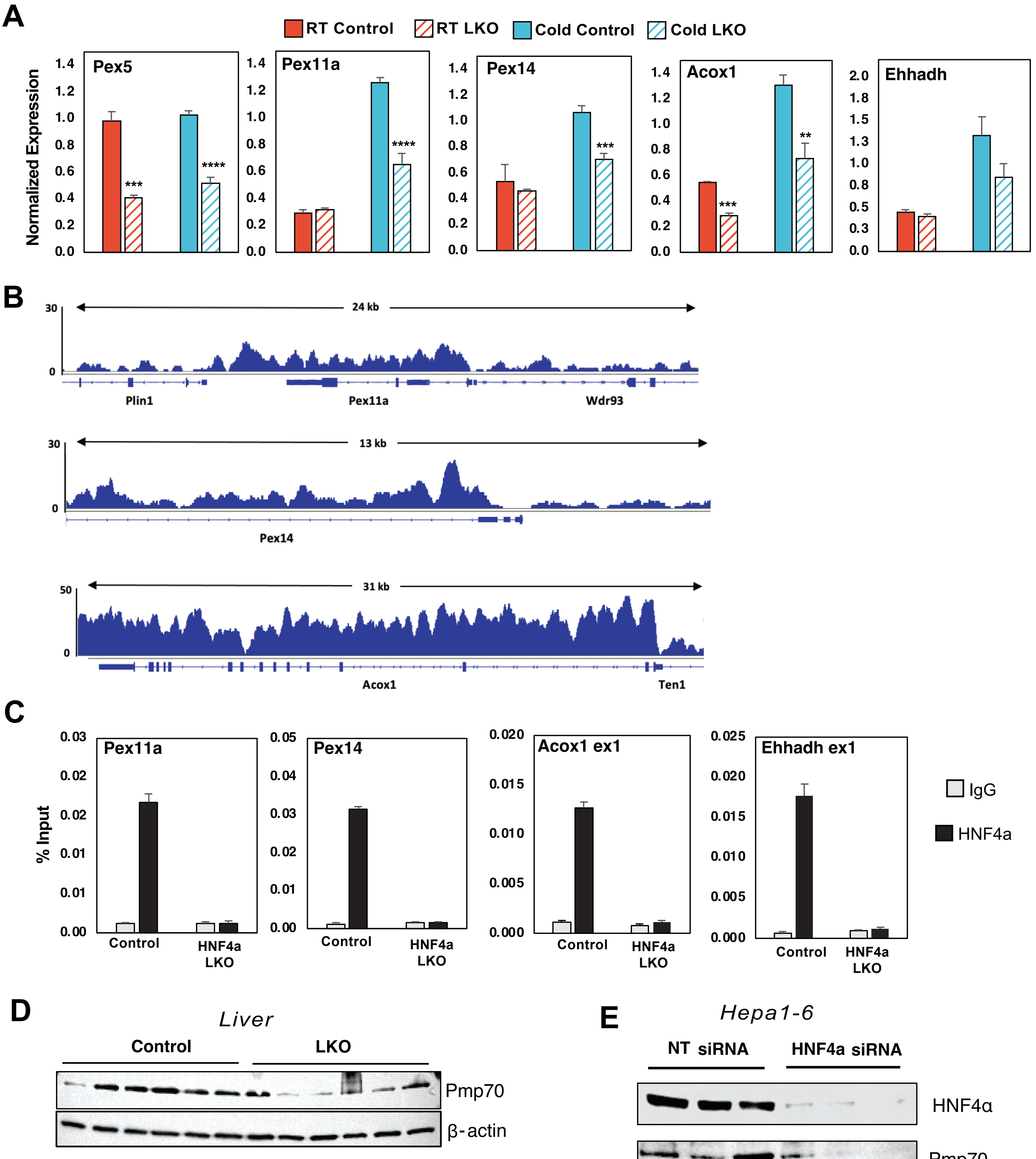
